# Towards a low CO_2_ emission building material employing bacterial metabolism in a two-step process of limestone dissolution and recrystallization: The bacterial system and prototype production

**DOI:** 10.1101/535161

**Authors:** Anja Røyne, Yi Jing Phua, Simone Balzer Le, Ina Grosås Eikjeland, Kjell Domaas Josefsen, Sidsel Markussen, Anders Myhr, Harald Throne-Holst, Pawel Sikorski, Alexander Wentzel

## Abstract

The production of concrete for construction purposes is a major source of anthropogenic CO_2_ emissions. One promising avenue towards a more sustainable construction industry is to make use of naturally occurring mineral-microbe interactions, such as microbial-induced carbonate precipitation (MICP), to produce solid materials. In this paper, we present a new process where calcium carbonate in the form of powdered limestone is transformed to a binder material (termed BioZEment) through microbial dissolution and recrystallization. For the dissolution step, a suitable bacterial strain, closely related to *Bacillus pumilus*, was isolated from soil near a limestone quarry. We show that this strain produces organic acids from glucose, inducing the dissolution of calcium carbonate in an aqueous slurry of powdered limestone. In the second step, the dissolved limestone solution is used as the calcium source for MICP in sand packed syringe moulds. The amounts of acid produced and calcium carbonate dissolved are shown to depend on the amount of available oxygen as well as the degree of mixing. Precipitation is induced through the pH increase caused by the hydrolysis of urea, mediated by the enzyme urease, which is produced *in situ* by the bacterium *Sporosarcina pasteurii* DSM33. The degree of successful consolidation of sand by BioZEment was found to depend on both the amount of urea and the amount of glucose available in the dissolution reaction.

## 1. Introduction

Globally, more than 10 km^3^ of concrete is produced every year [1], making it by far the most used construction material. Concretes are made from water, aggregates (coarse and fine particles such as gravel and sand), and cement that acts as a binder between the particles. A main ingredient in most common cements is lime (CaO), which is produced by high-temperature calcination of crushed limestone. Limestone is a sedimentary rock composed primarily of calcium carbonate (CaCO_3_), originating from chemical and biochemical processes that removed CO_2_ from the atmosphere in the geological past, often tens to hundreds of millions of years ago. The fossil CO_2_ released from limestone during calcination, and from burning of fossil fuels to drive this process, currently accounts for more than 5% of global anthropogenic CO_2_ emissions [2]. One possible avenue to significantly reduce the environmental footprint of the concrete industry is to employ naturally occurring mineral-microbe interactions in the production of construction materials [3-5] thereby replacing extensive heating requirements.

A number of approaches applying biotechnology in construction are currently pursued [5]. Most of them focus on biocementation through the concept of microbial-induced carbonate precipitation (MICP) [3, 4]. Several bacterial species can induce precipitation of calcium carbonate minerals in the presence of carbonate and calcium ions through metabolic processes that increase the pH in their surroundings. Most widely used is the production of the enzyme urease to hydrolyze added urea. This hydrolysis both increases pH through liberation of ammonia and generates carbonate. Research on MICP focuses on small-scale applications such as stone conservation [6, 7], crack remediation in concrete [8, 9], soil stabilization [10], self-healing concrete [11], and construction of subsurface barriers against pollutant spread or reservoir leakage [12].

However, large-scale replacement of conventional cement with a microbially produced binder in concrete still remains a considerable challenge [13]. Biocementation is a complex process as the microbial activity depends on a large number of environmental factors such as temperature and nutrients [14], and there is a lack of field tests and investments in large-scale geotechnical and industrial investigations [5]. Technical challenges include processing time [13, 14], the behavior of precipitated material from different reactants, which affects the porosity, the distribution of precipitated material, and the binding strength between the particles [4]. In the long term, chemical changes caused by the ammonium from urea hydrolysis may have adverse effects on the precipitated binder [15], and nutrient residues and metabolic by-products inside the material may lead to colonization by other microorganisms and affect the long-term performance of the material [4, 14].

A major challenge for large-scale biocementation is a suitable calcium source. Most commonly used is CaCl_2_, which is a by-product in the Solvay process for Na_2_CO_3_ production. However, the use of CaCl_2_ generates chloride ions that can drive corrosion of steel used as reinforcement [3]. Alternative calcium sources that have yielded promising results include calcium lactate [3], calcium hydroxide, calcium nitrate, calcium acetate, and calcium released from eggshells by treating them with acetic acid [16]. For all of these alternatives, cost and availability can limit large-scale application [3].

In this study, we introduce a new avenue for biocementation where powdered limestone, which is a cheap and abundant raw material, *e.g.* as waste fines from limestone quarries [16], is used as calcium source. It has recently been suggested by Choi *et al.* [16] that limestone can be dissolved by acetic acid, a by-product from biofuel production, to produce the Ca^2+^-containing solution required for MICP. The interplay between bacterially produced organic acid to dissolve calcium carbonate and subsequently precipitate it in the presence of urea has been described in natural environments [17, 18], but has not yet been applied in biocementation for construction purposes. In our study, we used acid producing bacteria to dissolve a dense slurry of powdered limestone as the first step in a two-stage process outlined in Figure 1. As acid is produced, the pH of the system drops, and Ca^2+^ ions are released. This process requires acid-producing bacterial strains that can grow and produce acid in contact with solid calcium carbonate at an initially high (around 9.5), but subsequently decreasing pH (down to approx. 6). The strains must tolerate a relatively high ionic strength, and require a minimum of growth supplements and medium components in order to limit both nutrient residues inside the final material, and the overall production costs. In addition, the strains must be non-pathogenic and easy to store and handle in large-scale manufacturing processes. In the next step, the solution containing high concentrations of free Ca^2+^ and bicarbonate ions is added to a packing of aggregate together with limited amounts of urea and urease producing bacteria. When urease is produced and hydrolyzes urea, pH increases and the dissolved limestone re-precipitates *in situ* as a calcium carbonate binder between the aggregate grains. We named this new binder BioZEment, with the capitals ZE representing the term ’Zero Emission’ to underline our efforts to produce a low CO_2_ emitting cement. The resulting material is similar to a porous sandstone.

**Figure 1:**
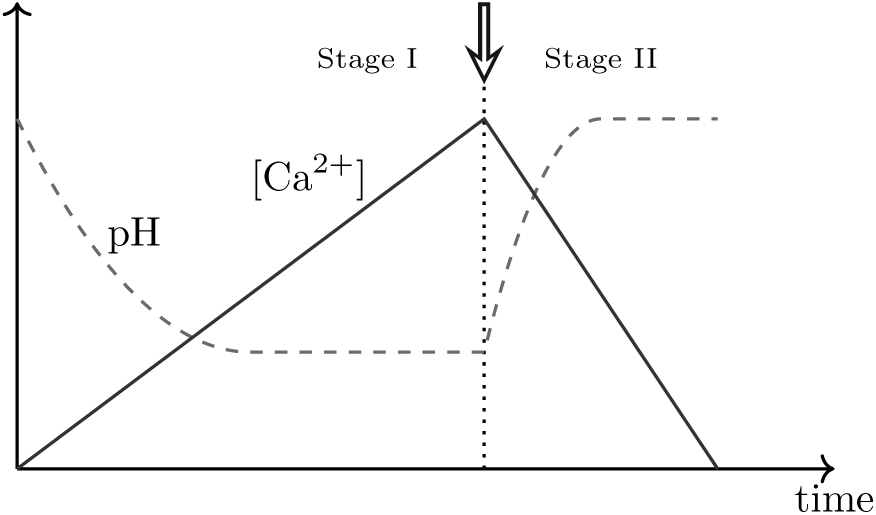
Schematic overview of the proposed two-stage limestone dissolution-recrystallization process. In Stage 1, bacteria produce acid in a dense slurry of calcium carbonate, reducing the pH, limestone dissolves, and the concentration of Ca2+ in solution increases. To initiate Stage 2, a second, urease-producing bacteria is added together with urea (indicated by the arrow). The urease hydrolyses the urea, causing the system pH to increase and Ca^2+^ and HCO_3_^-^to precipitate from the solution as calcite.

Life Cycle Assessment (LCA) and techno-economic analysis of a range of possible construction products using the proposed new method indicate that it can be a feasible alternative to a significant volume of conventional concrete, both in terms of cost and climate impact **(Myhr et al, submitted)**.

## 2. Materials and methods

We have performed three series of experiments, aimed at elucidating the general feasibility and current limitations of the proposed biocementation process (Figure 2).

**Figure 2:**
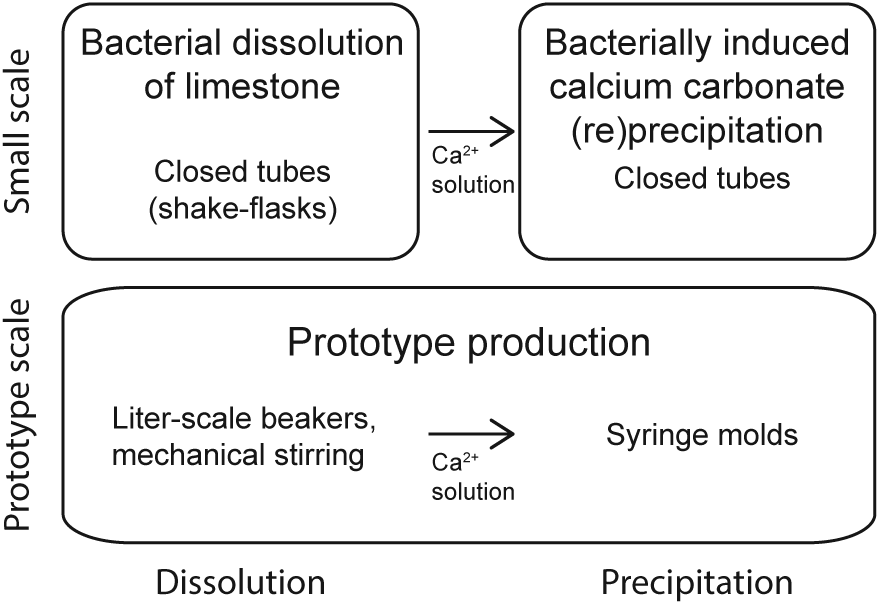
Experimental design of the present study. Small-scale experiments were carried out separately for Stage 1 and Stage 2 of the overall process depicted in Figure 1 in order to screen a number of variables and provide real-time monitoring of parameters such as pH and acid production. Parameters used in prototype production experiments were chosen based on results from the small scale dissolution and (re)precipitation experiments.

We first investigated the organic acid production and limestone dissolution in limestone slurry-bacteria systems under a range of processing parameters, including limited oxygen supply and a high initial pH, using the bacterial strain AP-004, isolated from a high-pH natural environment. Next, we studied the increase in pH and resulting decrease in Ca^2+^ concentrations in solution by adding various amounts of urea and the urease producing strain *Sporosarcina pasteurii* DSM33, a commonly used strains in MICP applications. Finally, we applied the most promising processing conditions to produce Ca^2+^-containing solutions at litre scale and used these to produce centimetre-sized consolidated samples as proof-of-concept of solid material generation and for material properties analysis.

### 2.1 Strains and cultivation conditions

#### Bacterial strains

The experiments were carried out using the acid-producing strain AP-004 (NCIMB accession number 42596), closely related to *Bacillus pumilus*, and the urease-producing strain *S. pasteurii* DSM33 obtained from the "Deutsche Sammlung von Mikroorganismen und Zellkulturen" (DSMZ). AP-004 originates from soil collected near an open CaCO_3_ quarry in Tromsdalen (Verdal, Norway) from which a heat-treated freeze-dried water extract was prepared. The strain was cultivated on rich medium, pH 9.5 (RM 9.5; see below), containing 10 g/L glucose. Strain identification was carried out using PCR (primer pair SD-bact-0011 5′-GTTTGATCCTGGCTCAG-3′ + SD-bact-1492 5′-ACGGYTACCTTGTTACGACTT-3′) to amplify the 16S rRNA gene followed by sequencing (Eurofins Genomics, Ebersberg, Germany).

#### Cultivation conditions

AP-004 was routinely cultivated in liquid or on solid rich medium, pH 9.5 (RM 9.5), consisting of, in g per litre; 1.0 yeast extract (Oxoid), 3.0 peptone (Oxoid), 7.5 glucose, 5.0 NaCl, 10.0 limestone powder (1-200 μm BETOFILL-Sa, Franzefoss Miljøkalk, Norway), and 1.2 Na_2_CO_3_. For solid medium, 15 g/L agar was added. To assess the microbial acid production visually as a colour change from blue to yellow, 2.5 mL/L thymol blue solution (10 g/L) was added. Deviations from standard glucose and limestone powder concentrations are indicated in the text. *S. pasteurii* was grown in DSMZ Medium 220 with 20 g/L urea.

#### Preparation of freeze-dried cells

To prepare a stock of freeze-dried cells, *S. pasteurii* was grown overnight, and centrifuged to remove the culture broth (2900 x g, 30 min). The cell pellet was washed once with sterile Milli-Q water and suspended in 100 g/L sucrose to a 7 times concentrated cell suspension. One mL aliquots of the cell-sucrose mixture were added to 10 mL sterile Hungate tubes covered with a paper cap and frozen at -80 °C. The frozen suspensions were freeze-dried at room temperature and less than 0.1 mbar for approx. 48 hours. The tubes containing dried cells were transferred to an anaerobic workstation where the paper cap was replaced by a sterile septum and a screw cap. The headspace of the Hungate tube was flushed with sterile-filtered nitrogen. Freeze-dried cells were stored at -20 °C.

### 2.2 Bacterial dissolution of limestone

AP-004 was plated onto RM 9.5 solid medium plates and incubated for two to three days at 30 °C prior to storage at 4 °C. Fresh RM 9.5 liquid medium was inoculated with cells from this agar plate and incubated at 30 °C with rotation for two to three days. Cells were pelleted by centrifugation (1, 600 x g, 15 min) and washed with fresh medium without glucose. Variants of RM 9.5 liquid medium were prepared with or without limestone powder (final concentration 50 g/L), and varying glucose concentrations, and inoculated with the washed cells from the AP-004 pre-culture. Cultivations were performed at ambient temperature (22-25 °C) in either 500 mL baffled shake flasks with 100 mL medium, or in 50 or 125 mL plastic tubes with 30 mL or 100 mL medium, respectively, and different agitation regimes representing different conditions for gas exchange with the environment. Samples (2.5 mL) of the resulting Dissolved Limestone Solution (DLS) were withdrawn at intervals for pH, Atomic Absorption Spectroscopy (AAS), and High-Performance Liquid Chromatography (HPLC) analysis (see below). Samples were centrifuged (18, 000 x g, 10 min), and the supernatants and pellets frozen separately at -20 °C until further analysis.

### 2.3 Bacterially induced calcium carbonate (re)precipitation

Fresh DSMZ Medium 220 (up to 60 mL) with 20 g/L urea was inoculated directly from a frozen *S. pasteurii* glycerol stock and incubated for about 24 hours at 30 °C with vigorous shaking. Bacteria were pelleted by centrifugation (1, 600 x g, 15 min) and washed once with peptone-saline (per litre: 1 g peptone, 8.5 g NaCl) or saline (per litre: 9 g NaCl). Bacterially dissolved limestone solution (DLS, 30 mL) (see 2.2) was mixed with a 2% (v/v) suspension of washed *S. pasteurii* cells and added urea to a final concentration of 20 g/L (unless otherwise stated) in 50 mL tubes, and incubated at room temperature. Controls without *S. pasteurii* were included, and experiments performed in triplicates. Samples (1.5 mL) for pH, AAS, and HPLC analyses were taken immediately before and after addition of *S. pasteurii* and urea, as well as at intervals during urea hydrolysis until pH remained constant.

### 2.4 Prototype production

RM 9.5 medium (50 mL) was inoculated with AP-004 cells from an agar plate (stored at 4 °C) and incubated with shaking at room temperature for at least 2 days. 40 or 80 mL of the pre-culture was added directly to 1 or 2 litres RM 9.5 medium containing 40 or 80 g limestone powder, respectively, and with glucose concentrations of 7.5, 10, or 30 g per litre (Table 1). The solution was stirred continuously using a mechanical stirrer (VWR VOS 14, anchor stirrer, 500 rpm) for 2-3 days at room temperature. The pH of the solutions was measured directly after stirring. The solution was filtered (11 µm pore size filter paper) to remove remaining limestone particles. In some experiments, the bacteria were also removed by micro-filtration (Millipore Stericup filters, pore size 0.22 μm). The resulting solutions were stored at 4 °C for up to two weeks. Samples for AAS and HPLC analysis were taken of the freshly prepared solutions, as well as at the end of storage time. Samples for AAS and HPLC were centrifuged (10, 000 x g, 10 min), and the supernatant frozen at -20 °C until analysis.

**Table 1:** Overview of experimental parameters used in prototype sample production. DLS: Dissolved limestone solution.

| Sample no. | Initial glucose conc. in DLS (g/L) | Filtered (F)/ non-filtered (NF) DLS | Urea (g/L) | Limestone powder (g/kg, total solids) | Number of replicates | Injection direction |
| --- | --- | --- | --- | --- | --- | --- |
| 1 | 30 | F | 1.5 | - | 3 | up |
| 2 | 30 | F | 3.0 | - | 3 | up |
| 3 | 30 | NF | 6.0 | - | 3 | up |
| 4 | 30 | NF | 1.5 | - | 3 | up |
| 5 | 30 | NF | 3.0 | - | 3 | up |
| 6 | 30 | NF | 6.0 | 25 | 3 | up |
| 7 | 30 | NF | 6.0 | 50 | 3 | up |
| 8 | 30 | NF | 3.0 | 25 | 3 | up |
| 9 | 30 | NF | 3.0 | 50 | 3 | up |
| 10 | 30 | NF | 1.5 | 25 | 3 | up |
| 11 | 30 | NF | 1.5 | 50 | 3 | up |
| 12 | 30 | F | 40 | - | 3 | up |
| 13 | 10 | F | 40 | - | 3 | up |
| 14 | 30 | NF | 40 | - | 3 | up |
| 15 | 7.5 | NF | 8.0 | - | 3 | up |
| 16 | 7.5 | NF | 40 | - | 3 | up |
| 17 | 10 | NF | 8.0 | - | 3 | up |
| 18 | 10 | NF | 40 | - | 3 | 2 up, 1<br>down |
| 19 | 7.5 | NF | 8.0 | 25 | 3 | down |
| 20 | 7.5 | NF | 40 | 25 | 3 | down |
| 21 | 10 | NF | 8.0 | 25 | 3 | down |
| 22 | 10 | NF | 40 | 25 | 3 | down |
| 23 | 30 | NF | 8.0 | 25 | 3 | down |
| 24 | 7.5 | NF | 6.0 | 25 | 1 | down |
| 25 | 30 | NF | 6.0 | 25 | 1 | down |

A stock solution of *S. pasteurii* DSM33 was prepared by inoculating 50 mL DSMZ medium 220 with frozen, lyophilized culture and incubation on a shaking table overnight at ambient temperature (22-25 °C). From this culture, 1 mL was used to inoculate 50 mL fresh DSMZ medium 220 and incubated under the same conditions for one day, before storing the stock solution at 4 °C for up to two weeks. The injection reagent for prototype production was prepared by mixing DLS from prototype-scale dissolution as described above with the stored *S. pasteurii* culture, diluted 50 times in DSMZ medium 220, and various amounts of urea stock solution (400 g/L) resulting in final urea concentrations ranging from 1.5 to 40 g/L. Reagents with different initial amounts of glucose in the DLS and urea were prepared as presented in Table 1.

Prototype samples were produced by 40 injections of injection reagent into 20 ml syringes (HSW SOFT-JECT, 20 mL, inner diameter 20 mm), filled with sand (50-70 mesh size from Honeywell Specialty Chemicals Seelze GmbH, Germany). In some samples (Table 1), the sand was mixed with limestone powder. The inside of the syringe was sprayed with Teflon (Norton SprayFlon, DuPont) to facilitate recovery of the sample from the mould after consolidation. A 5-mm thick 3D printed plastic drainage grate and an 11 µm pore size filter paper were placed at the bottom of the syringe, and another filter paper placed on top. The sand and limestone powder were mixed by manual stirring, poured loosely into the syringe, and compacted by pressing down a rubber plug, which was also used to seal the system, onto the top filter paper. The experimental setup is shown in Figure 2.

Fluid injections were performed manually with 20 mL syringes connected to the prototype moulds via silicon tubes (3 mm inner diameter). In some cases (Table 1), the liquid was introduced from the bottom and drained via a silicon tube through the rubber plug at the top of the syringe (Figure 3). To facilitate simultaneous injection to several samples, a rig was constructed where the plungers of the injection syringes for samples 18-25 were all simultaneously pressed down by one large plate. In this setup, the injections were done from the top, through the rubber plug, and the fluid drained from the bottom. Every injection consisted of 12 mL reagent, which was sufficient to completely replace the fluid in the sample. Between the injections, the injecting syringes remained attached to the inlet tubes so that the system was only exposed to the atmosphere via the exit of the drainage tube. To prevent spilling of reactants, the inlet tubes were filled with about 2 mL of air at the end of each injection, pushing all the injected reactants into the sample. This air was subsequently injected into the sample at the start of the next injection. The minimum interval between injection was 5 hours with no more than two injections per day. The total number of injections was 40, thus one production experiment was completed in approx. 4 weeks. Samples of the effluent were taken once every 10 injections and frozen at -20 °C until AAS analysis.

**Figure 3:**
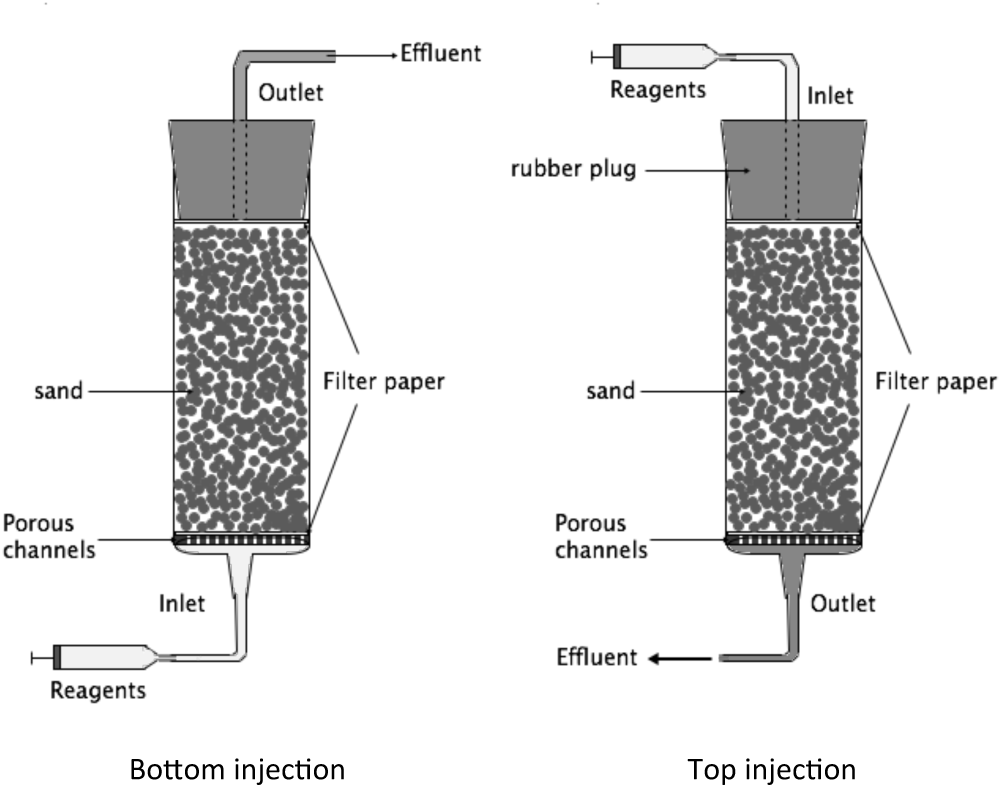
Experimental setup for prototype production. A 20 ml syringe was used as a mould. A drainage grate with a filter paper on top was placed at the bottom of the syringe. Next, the syringe was filled with sand, a filter paper placed on top of the sand, and the sand somewhat compacted with the top rubber plug. Bottom injection (left): Liquid was introduced from the bottom through a silicon tube and drained through the rubber plug at the top. Top injection (right): Liquid was introduced from the top in a reciprocal setup.

After 40 injections, the samples were rinsed with two full injections of MilliQ water, and, while still inside the moulds, dried overnight at 60 °C, before finally removed from the moulds and dried overnight at 90 °C.

### 2.4 Chemical analysis

#### pH

In limestone dissolution experiments, pH was monitored directly in the limestone dissolving culture using a Radiometer Analytical PHC3359-8 combination pH electrode for microsamples (Hach) calibrated with two standard solutions. In (re-)precipitation experiments, pH was measured at the end of the dissolution experiments using an ELIT P2011 combination pH electrode (Nico2000), which was calibrated using three standard solutions.

#### HPLC

Samples were thawed at room temperature and filtered (0.2 µm) to remove any particles. Glucose and organic acid concentrations were determined by HPLC using either a Shimadzu Model 9A instrument or an Agilent HPLC 1600 system with a DAB-UV-and an RI detector, and equipped with an Aminex HPX-87H column (Bio-Rad). Compounds were eluted with 5 mM H_2_SO_4_ at a flowrate of 0.6 ml/min and 45 °C. Quantification was performed using standards of lithium lactate, sodium acetate, sodium pyruvate, sodium propionate, sodium succinate, sodium citrate, and glucose with known concentrations.

#### LC-UV-QTOF

Unknown organic compounds were identified by mass spectrometry. High resolution LC-MS analysis was performed on an Agilent 1200 Series LC/MS system (Agilent Technologies Inc., Palo Alto, CA, USA) connected to an Agilent 6530 QTOF mass spectrometer. LC separation was performed in RP mode employing a Zorbax XDB-C18 (4.6 x 50mm, 3.5 µm particle size) column and gradient elution using 25 mM formic acid as Eluent A and acetonitrile as Eluent B as follows: idle at 5% B, 1 min 5% B, 4 min ramp to 50% B, 5 min hold at 50% B, 30 s ramp to 5%. The flow rate was 0.3 ml/min, and the column thermostat was maintained at 30 °C. The injection volume was 5 µL. All analyses were performed with electrospray ionization (ESI) in either positive or negative scan mode. The mass range was 50 to 2500 Da. The gas temperature was 350 °C, and the gas flow was 9 L/min. The sheath gas settings were 10 L/min and 350 °C. The nebulizer was set to 45 psig. The capillary voltage was set to 3500 V, the nozzle voltage to 1000 V, and the fragmentor to 150 V. In addition to the mass spectrometry detection, the UV absorbance was monitored from 190 to 640 nm. For each sample, a list of possible compound masses was generated, and the most abundant compound masses were compared to the masses present in the media corresponding to each sample. The UV chromatograms for the samples and the media samples were also compared, and the mass spectra corresponding to UV signals present in the samples but not in the media were collected.

#### AAS

The total concentration of dissolved calcium was quantified using atomic adsorption spectroscopy (AAS) with a Perkin Elmer AANALYST400.

### 2.5 Material characterization

#### Compressive strength

Consolidated samples were capped with gypsum on both ends to ensure that the test cylinders had smooth, parallel, uniform bearing surfaces perpendicular to the applied axial load during the compression test. The specimens were subjected to uniaxial compression using an Instron 3345 universal testing machine (Instron, USA) with a 5 kN load cell and a cross-head speed of 10 mm/min.

#### XRD

Crystals present in the consolidated samples were identified using Rigaku MiniFlex600 X-Ray diffractometry, with a scan range from 10° to 90° and a 10°/min scanning rate. The X-Ray source provided Cu-Kα radiation with a wavelength of 0.154 nm.

#### SEM

Small pieces of the consolidation samples were imaged using a TM3000 table-top microscope (bench-top SEM) from Hitachi High-Technologies.

#### CaCO_3_ content measurement

Portions of the consolidated samples (0.4-1.0 g) were dried at 70 °C for 24 hours. Dried samples were added to 30 mL 1 M hydrochloric acid (HCl) in closed tubes and left for at least 6 hours on a shaking table in addition to repeated manual shakings of the tubes for a total of 48 hours at room temperature. The remaining solids were rinsed several times with DI water, followed by drying at 70 °C for 24 hours and re-weighing. The CaCO_3_ content was determined as the ratio of the sample weight before and after acid dissolution. In samples with added limestone powder, the amount of precipitated CaCO_3_ was determined by subtracting the initial CaCO_3_ concentration.

### 2.6 Thermodynamic calculations

Thermodynamic calculations were performed using the PHREEQC geochemical software [19] and the PHREEQC database. Data for calcium lactate species were taken from Vavrusova *et al.* [20]. Calculations were made for both, open systems, with the fluid in equilibrium with an atmospheric *p*CO2 of 10^-3.5^ atm, and closed systems without CO_2_ exchange with the atmosphere. However, for the closed system, the maximum *p*CO2 was set to 0.1 atm, which was assumed to be a reasonable upper limit for dissolved CO_2_ without gas exchange through bubble formation. Calculations for both system types were made assuming constant (atmospheric) pressure and a temperature of 20.0 °C.

## 3. Results

### 3.1 AP-004 produces acids from glucose in the presence of high concentrations of CaCO_**3**_**, providing sufficient pH reduction to dissolve limestone.**

For Stage 1 of the two-stage process outlined in Figure 1, a bacterial strain capable of producing acid at pH 6.0-9.5, and in the presence of increasing amounts of calcium and carbonate ions during the dissolution process was required. To identify a suitable strain, soil samples were collected in a natural environment with high limestone content and thus alkaline pH in the required range. After enrichment and isolation, the most suitable strain for proof-of-concept of the envisioned limestone dissolution process was chosen, a strain denoted AP-004. AP-004 was, based on 16S rRNA gene amplicon sequencing, found to be closely related to *Bacillus safensis* strains FO-36b and NBRC 100820 (100.00 % sequence identity based on the part of the 16S rRNA gene sequence covered). It showed robust and reproducible growth and acid production on RM 9.5 liquid medium both in the presence and absence of limestone powder.

Cultivations of strain AP-004 were performed in closed tubes to mimic limited oxygen availability as expected in a scalable dissolution process. Culture growth and acid production was observed both in the absence and presence of limestone powder (Figure 4). Culture growth in the absence of limestone (Figure 4A) started immediately after inoculation and reached a maximum OD_660_ of 0.8 approx. 13 days after inoculation, concurrent with a rapid drop in pH by more than two units within the first 24 h and leveling out at a pH below 6.0 at which culture growth ceased. In the presence of 10 g/L limestone powder (Figure 4B), it was not possible to follow culture growth as optical density of the culture. However, based on the observed similar decrease in pH under these conditions and a similar, constant rate of glucose consumption during the first 140 h compared to the culture without limestone, it appeared that culture growth proceeded in a similar manner. The pH after 140 h in the presence of limestone powder was slightly higher than in its absence, presumably due to the buffering effect of calcium carbonate. After 140 h, the glucose consumption rate in the presence of limestone was approximately twice the rate observed in its absence, suggesting that the low pH in the unbuffered system had an adverse effect on cell metabolism.

**Figure 4:**
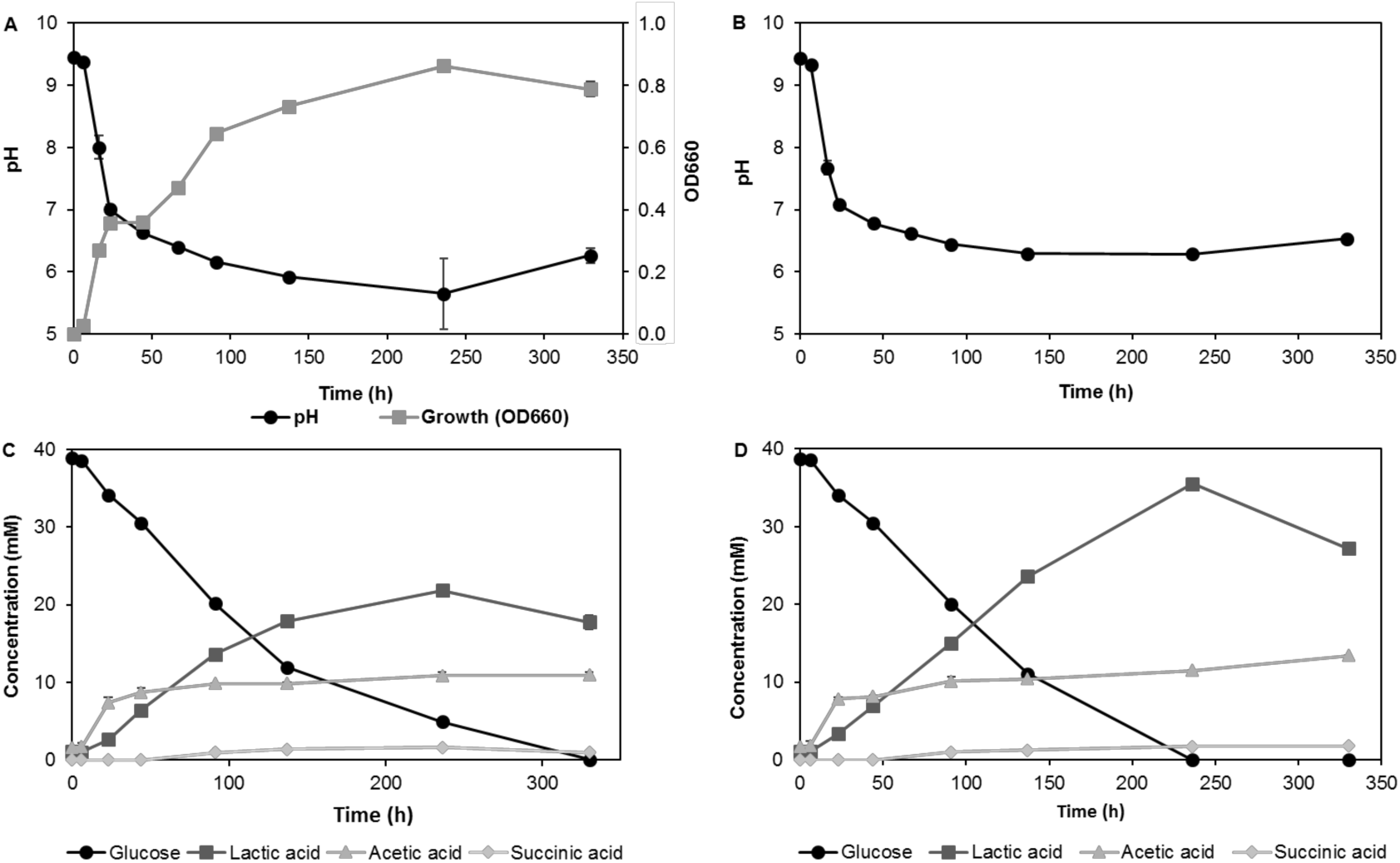
Cell growth, pH, glucose consumption, and acid production profiles of AP-004 cultures in the absence (A/C) and presence (B/D) of 10 g/L limestone powder. Error bars represent standard deviations between three independent cultures. AP-004 was cultivated in closed tubes containing RM, pH 9.5 at 30 °C.

The main organic acids produced under these conditions were lactic acid and acetic acid as confirmed by LC-MS. After 250 hours of incubation, lactic acid constituted more than 50% and 70% of the total organic acid (defined as the sum of lactic and acetic acid from here on) produced in the absence (Figure 4C) and presence (Figure 4D) of limestone powder, respectively. The initial 40 mM glucose was completely metabolized after 330 h and 240 h, respectively. The final concentration of total acids produced by the bacteria was at least 33 mM and 47 mM in the absence and presence of limestone powder, respectively. Both, the glucose consumption and the acid production rates in the period after the main growth phase (up to ca. 140 h) appeared to be higher in the presence of limestone powder, indicating that a pH below approx. 6.0 inhibited growth and acid production by AP-004. The consistent decrease in lactic acid concentrations in the final phase of the experiments (after ca. 240 h) and corresponding pH increase, may indicate that lactic acid was metabolized after depletion of the primary carbon source, glucose.

From the presented experiments, it can be concluded that the selected strain AP-004 can produce acids from glucose in the presence of high concentrations of limestone powder and limited oxygen supply, and that the acid production is sufficient to reduce the pH in the presence of limestone to a degree that enables its dissolution.

### 3.2 Thermodynamic modelling and experimental testing demonstrate that the system chosen is closed, CO_**2**_ **is largely contained, and pH drop and limestone dissolution proceeds independent of the type of acid produced.**

Thermodynamic calculations were performed to predict how the amount of produced acids and the amount of dissolved carbonate in the system influenced the solubility of CaCO_3_ (Figure 5). The concentration of carbonate species in solution are controlled by the following reactions [21]:

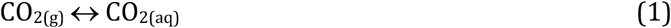

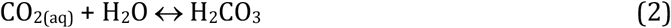

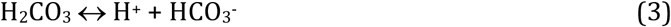

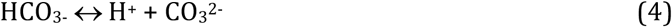

**Figure 5:**
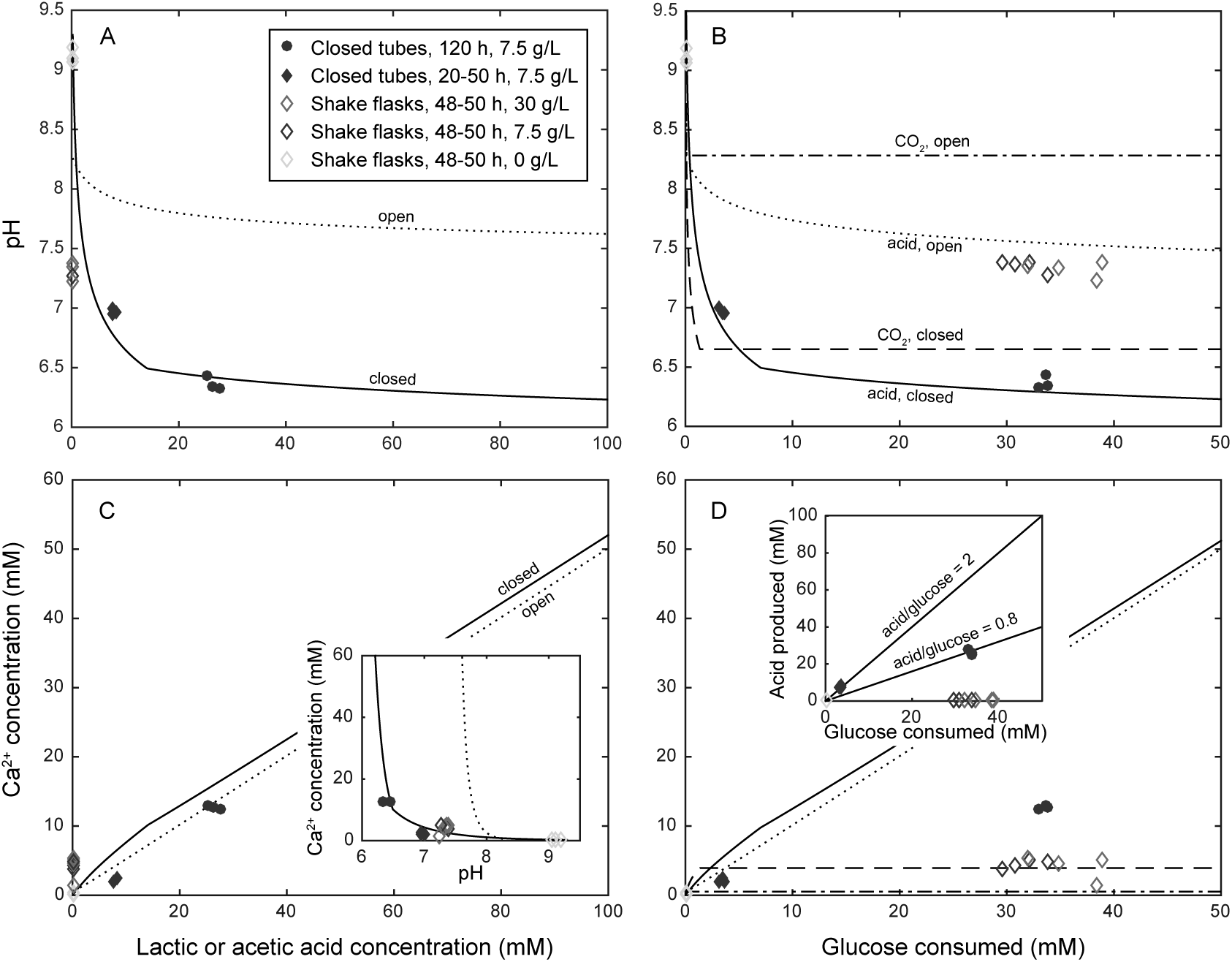
Results from thermodynamic calculations, showing pH (A and B) and total dissolved calcium carbonate (C and D) as a function of the concentration of produced lactic acid (A and C) and consumed glucose (B and D) for open and closed systems. The inset in panel C shows the same data plotted as Ca2+ in solution as a function of pH. Experimental data shown are for glucose levels of 30 g/L (red), 7.5 g/L (blue), and 0 g/L (green). Experimental conditions: closed tubes, 20-50 h dissolution (filled diamonds); closed tubes, 120 h dissolution (filled circles); shake flasks, 48-50 h dissolution (open diamonds). Calculated data presented in panels B and D are based on the assumption that all glucose is converted to acid (solid/dotted lines) or CO_2_ (dashed lines). The inset in panel D shows total amount of acid produced as a function of glucose consumed, where solid lines correspond to 2 and 0.8 moles acid per mole glucose, respectively.

At pH 6.3 to 10.3, coincidental with our experiments, HCO_3_^-^is the predominant carbonate species in solution. The solubility of CaCO_3_ is very low (K_sp_ = 3.3 · 10^-9^ [21]), but is dissolved by addition of acid:

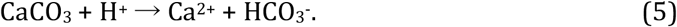

The produced HCO_3^-^_ may through reactions (1-3) be converted to CO_2(g)_. If the system is in equilibrium with atmospheric CO_2_ in air (’open system’), the low partial pressure of CO_2_ in air (approx. 0.0004 atm) means that some of the produced carbonate will escape from the system. However, although the transformation from dissolved to gaseous CO_2_ described by reactions (1) and (2) can occur through bubble formation throughout the system, provided the carbonate concentration is above a critical value for the given conditions, the gaseous CO_2_ can only escape from the system at the fluid-air interface. The rate of carbonate loss from the system is therefore limited by transport through the bulk fluid. Systems without vigorous shaking or stirring may therefore be considered as closed, with the dissolved carbonate contained in solution on the timescale of hours to days. The result is that the system pH will be lower than in the open system.

When the concentration of CO_2(aq)_ reaches a critical value, bubbles of CO_2(g)_ will form and as these bubbles grow, this will significantly enhance the rate of CO_2_ escape. Larger bubbles will move faster upwards in the system. This places an upper limit on the amount of carbonate that can be contained in the system. The critical concentration of CO_2(aq)_ depends on a range of parameters, such as the pressure and the presence of favourable bubble nucleation sites. We have chosen to model closed systems by limiting the concentration of CO_2(aq)_ to a level where it is in equilibrium with an atmospheric *p*CO2 of 0.1 atm (see Section 2.6).

According to the results presented in Section 3.1, lactic and acetic acid were the main organic acids produced by AP-004. The pK_a_ values for these two acids (3.86 and 4.76) are both well below the relevant pH range (6-10), and the calculations gave nearly identical results for the two acids. Thus, the system can be discussed in terms of total amount of acid produced without paying specific attention to the ratio between lactic and acetic acid. The results shown in Figure 5 are for lactic acid only.

As expected, the amount of dissolved carbonate has a large effect on the system pH for a given amount of produced acid (Figure 5A). In the open system, pH remains above 7.5 for 150 mM acid produced, whereas in the closed system, the pH at 150 mM acid is below 6.5. This shows that the pH in a system where bacteria produce acid and calcium carbonate dissolves will be significantly influenced by the factors that control the release of CO_2_ to the atmosphere, such as the degree of shaking or stirring. On the other hand, the solubility of CaCO_3_ is almost unaffected by the carbonate concentration and increases close to linearly with the amount of acid produced (Figure 5B).

In the dissolution experiments performed in closed tubes, measured values of pH as a function of acid concentration as determined by HPLC (Section 3.1) were found to fit well with the predictions based on models for closed systems (Figure 5A and 5B), showing that these systems were indeed closed and that CO_2_ was largely contained as bicarbonate ions in solution.

Based on stoichiometry, up to 2 moles lactic acid, 4 moles acetic acid, or 6 moles CO_2_ can be produced from 1 mole glucose. The ratio between produced acid and CO_2_ will primarily depend on the availability of oxygen, i.e. aeration. Good access to oxygen will promote metabolism of glucose to CO_2_. The decrease in pH caused by CO_2_ production in a closed system (Figure 5C) will lead to dissolution of about 3 mM CaCO_3_ (Figure 5D) through the overall reaction

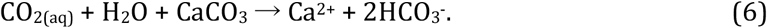

In a real system, some of the metabolized glucose will be converted to and remain bound in cell mass. The acid-to-glucose ratios of the experimental results shown in Figure 5D (inset) are between 2 and 0.8.

When dissolution experiments (AP-004, 7.5 g/L glucose in RM 9.5 liquid medium) were performed in an open system with good aeration (shake-flasks), lactic or acetic acid concentrations were below the detection limits of 0.67 mM and 1.34 mM, respectively (Figure 5A and 5C). Still, pH decreased to approx. pH 7.2 and around 3 mM Ca^2+^ was dissolved, confirming that CO_2_ produced by the cells plays a role in the overall acid balance in the system. However, the amount of calcium carbonate that can be dissolved by CO_2_ generation alone is very small compared to the amount that is dissolved when acid is produced, and 3 mM Ca^2+^ is most certainly insufficient for cementation in Step 2. This highlights the importance of limited oxygen supply for sufficient CaCO_3_ dissolution.

### 3.3 Addition of urea and urease producing *S. pasteurii* to bacterially dissolved limestone solution rapidly increases pH and precipitates CaCO_3_

*S. pasteurii* produces the enzyme urease that hydrolyzes urea (ureolysis) according to the reaction

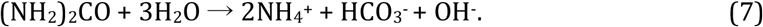

The formation of ammonium and hydroxide ions increases pH and induces calcium carbonate precipitation as required for Stage 2 in the process depicted in Figure 1:

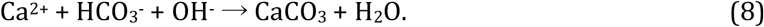

To demonstrate this effect, after culturing of AP-004 in RM 9.5 medium containing limestone powder for 120 h, variable amounts of *S. pasteurii* cells and a fixed amount of urea (20 g/L) were added to the bacterial culture. The results are presented in Figure 6. In these experiments, an initial glucose concentration of 7.5 g/L was used, as this amount of glucose had previously been found to be almost completely consumed within 120 h with a resulting pH below 6.5 and Ca^2+^-concentration above 10 mM (Figure 5).

**Figure 6:**
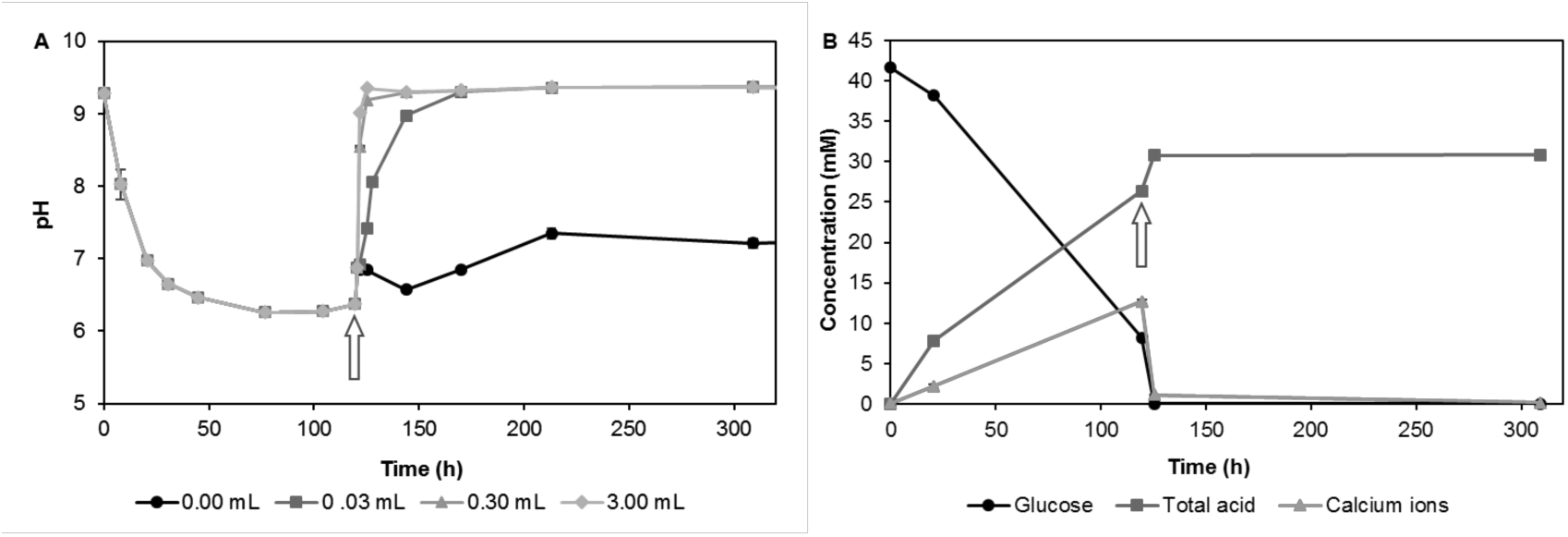
Results from experiments where AP-004 was cultivated in closed 50 mL tubes at room temperature with 7.5 g/L glucose (42 mM) from the start. After 120 h, 0-3 mL *S. pasteurii* culture and 20 g/L urea was added (marked with an arrow). A) pH development as a function of the amount of *S. pasteurii* culture added; B) acid production and released calcium ions in cultures in relation to glucose consumption for the DLS to which 0.03 mL *S. pasteurii* culture was added. The term "total acids" refers to the sum of lactic and acetic acid as determined by HPLC. The results are averages from at least two replicates.

The pH increased rapidly within a few hours after addition of *S. pasteurii* and urea, and the rate of pH change increased with the amounts of cells added. As an example, after 5 hours, the pH increased from 6.4 to 9.4 after addition of 3 mL *S. pasteurii* and urea. A control with only urea, but no *S. pasteurii* added, resulted in only a slight increase in pH (from 6.5 to 7), possibly because lactic acid was metabolized after all the glucose had been consumed (see above), small amounts of free ammonia in the urea, or chemical or biological hydrolysis of some of the added urea. Alternatively, AP-004 may, after the glucose had been depleted, have started to metabolize some of the amino acids present in the medium, leading to the observed pH increase. AP-004 grows on complex compounds in the absence of glucose (data not shown). The concentration of calcium ions dropped rapidly after addition of *S. pasteurii* and urea, indicating that CaCO_3_ precipitated due to the reduced solubility at high pH and the presence of carbonate ions.

The results revealed that the second stage of the process proceeded much faster than the acid dissolution in Stage 1, even when only small amounts of *S. pasteurii* were added. However, the presence of *S. pasteurii* cells were necessary to rapidly increase pH above 9. As ureolysis proceeded rapidly at a stage when almost all glucose had been consumed, obviously only minute amounts of glucose are needed to initiate and complete Stage 2.

With an initial concentration of 7.5 g/L glucose in RM 9.5 liquid medium, glucose was completely consumed after 125 h when no or only a very small amount (0.03 mL) of *S. pasteurii* culture was added. However, if 10-or 100-fold more *S. pasteurii* culture was added, consumption of the added glucose proceeded at a reduced rate. While glucose was completely consumed five hours after addition of 0.03 mL *S. pasteurii* culture, 1.6 and 3 mM glucose were detected in tubes to which 0.3 and 3 mL had been added, respectively. Even at the last sampling point (309 h), the glucose concentration was still around 0.7 mM. The slower rate of glucose consumption may be a result of a slower metabolism of AP-004 in response to the sudden increase in pH or by nutrient mass transfer reduction by the cells potentially being encapsulated by precipitated CaCO_3_. Even after more than 300 hours, lactate and acetate were still present, indicating that neither AP-004 nor *S. pasteurii* metabolized these compounds in significant amounts under the given conditions (Figure 6B). Further studies are necessary to elucidate if this apparent contradiction to results presented above (Figure 4) is solely an effect of the different pH’s in the two different situations or if other, yet unknown factors contribute.

### 3.4 A small-scale mould injection strategy yielded solid, stable material specimens in which newly formed calcite crystals efficiently bind aggregate

After demonstrating viability of the two-stage process and defining suitable conditions for both limestone dissolution and subsequent urease induced pH increase to trigger CaCO_3_ recrystallization, the concept was demonstrated by employing the consolidation process illustrated in Figure 1 to produce consolidated sand samples. Table 1 summarizes the parameters used in the consolidation experiments using BioZEment.

DLS was prepared in batches of 1 or 2 liters, with continuous mechanical stirring, with 7.5, 10, or 30 g/L glucose in the initial solution. Dissolution proceeded at room temperature for 2-3 days. For most of the samples, the DLS was only roughly filtered to remove undissolved limestone, while AP-004 cells could still be present in solution. Other samples were in addition sterile filtered in order to rempve the AP-004 cells. After filtering, the DLS was stored at 4 °C for up to two weeks during prototype production. No significant changes in dissolved organic compounds in the DLS were observed during storage.

HPLC analyses of the resulting DLS showed that added glucose was completely consumed when 7.5 and 10 g/L glucose was added from start, while 40-50% of the initial glucose still remained after 3 days for 30 g/L initial concentration. Lactic-and acetic acid were the most abundant metabolic products. Small amounts of succinate and pyruvate were also detected in some samples. The final concentrations of total acid were 65 ± 8 mM, 86 ± 13 Mm, and 180 ± 42 mM for initial glucose concentrations of 7.5 g/L (42 mM), 10 g/L (56 mM), and 30 g/L (170 mM), respectively, corresponding to molar acid-to-glucose ratios of 1.55, 1.54, and 1.78. The results indicate that most of the glucose was metabolized to acid, and not CO_2_, as a consequence of limited aeration (Figure 5D). The glucose consumption rate in these experiments was faster than those observed in closed tube experiments (Section 3.1, Figure 4**Error! Reference source not found.**), probably due to better mixing using a mechanical stirrer.

Syringes filled with packed aggregates (sand with or without powdered limestone, 25 or 50 g limestone/kg total solid) were injected with a mixture of DLS, urea (concentrations of 1.5 – 40 g/L) and *S. pasteurii* culture 40 times before rinsing with clean water, drying, removal from the moulds and final drying. Visual inspection (Figure 7) revealed that some of the processing parameters produced solid, consolidated samples, while others yielded little or no cohesion between aggregate grains. The quality of cementation increased with increasing urea concentration. Samples with less than 6 g/L (100 mM) urea did not cement well, and 40 g/L (666 mM) urea yielded better results than 8 g/L (133 mM) urea.

**Figure 7:**
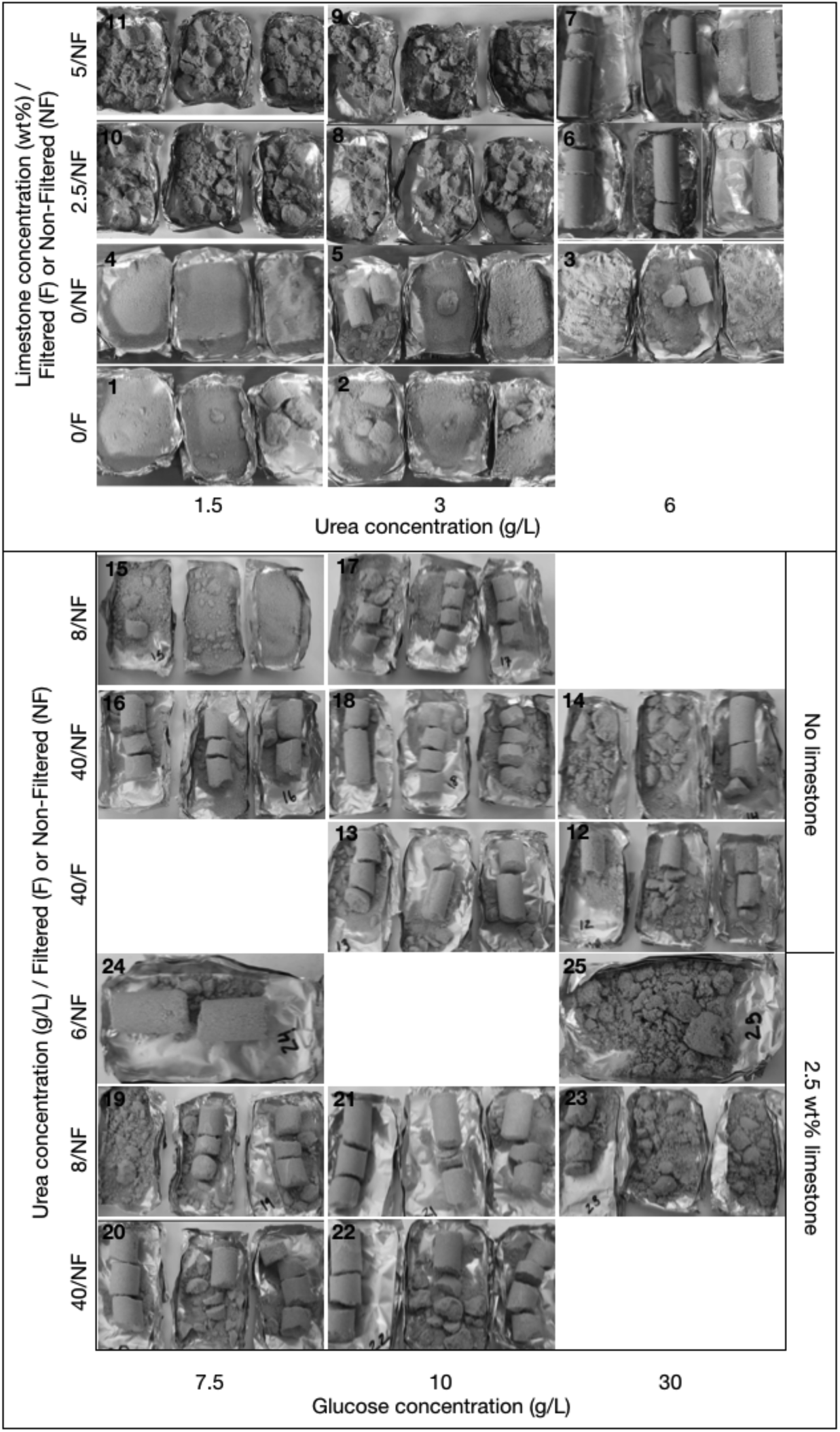
Visual results of prototype production. Numbers in the top left corner of each image refers to sample numbers (Table 1).

Samples containing limestone powder showed better consolidation than those without, which may be due to the difference in grain size distribution and/or the surface properties of calcium carbonate.

Comparing initial glucose concentrations in the preparation of DLS, the intermediate concentration (10 g/L) yielded the best results. According to the amounts of acid produced, there should be a higher Ca^2+^ concentration in the 30 g/L glucose DLS, which should again lead to a larger amount of precipitated CaCO_3_. AAS analyses of waste solutions after every 10^th^ injection showed that the concentration of Ca^2+^ was low, and decreased with each injection for the 7.5 and 10 g/L glucose derived DLS, indicative of continued urease production by *S. pasteurii*. For 30 g/L, the Ca^2+^ concentration in waste solutions remained very high for both filtered and unfiltered DLS. As there was also no significant difference in consolidation between filtered and non-filtered solutions, the presence of AP-004, although potentially capable of producing acid from glucose inside the material, is probably not the cause of the inferior quality of cementation for 30 g/L glucose derived DLS. This suggests that the presence of glucose somehow adversely affected the precipitation of calcium carbonate in this system, either indirectly by negatively affecting urease production by *S. pasteurii*, or directly by glucose or a derived metabolic product inhibiting calcite precipitation. Further investigations are needed to elucidate this effect in more detail.

XRD spectra (Figure 8) show the presence of quartz and traces of orthoclase feldspar in the sand, along with precipitated calcium carbonate in the form of calcite. No peaks corresponding to vaterite or aragonite, which are the other calcium carbonate phases that might be expected in this system, were found. SEM observations (Figure 9) showed that the precipitated calcium carbonate formed a coating on the aggregate grains, with individual calcite crystals of less than 10 μm size. Sites where aggregate grains had fallen off during sample preparation (see 2/NF/10), displayed a smooth surface of calcium carbonate that would originally have formed a close fit to the sand surface. Samples based on 10 g/L glucose derived DLS preparations had denser layers of crystals than samples based on 7.5 and 30 g/L glucose derived DLS preparations. Only in the sample based on filtered DLS preparation from 30 g/L initial glucose concentration, did some of the calcite form larger, spherical crystals typical of calcite precipitation with low nucleation rate in the presence of organic compounds such as lactate [22]. Because Ca^2+^ has a tendency to bind to the negatively charged surfaces of bacterial cells, bacteria of several species can act as favourable nucleation points for calcium carbonate precipitation and thus accelerate precipitation of CaCO_3_ [23]. It is therefore reasonable to assume that the use of micro-filtered DLS, where the AP-004 cells had been removed prior to injection, resulted in a lower nucleation rate. In samples added limestone powder, the calcite formed a less uniform coating on the sand grains than in the systems without limestone, indicating that more of the calcite precipitation occurred on limestone particles, where calcite nucleation is more favourable [6]. Bacterial attachment is also expected to be enhanced on limestone relative to sand surfaces [6].

**Figure 8:**
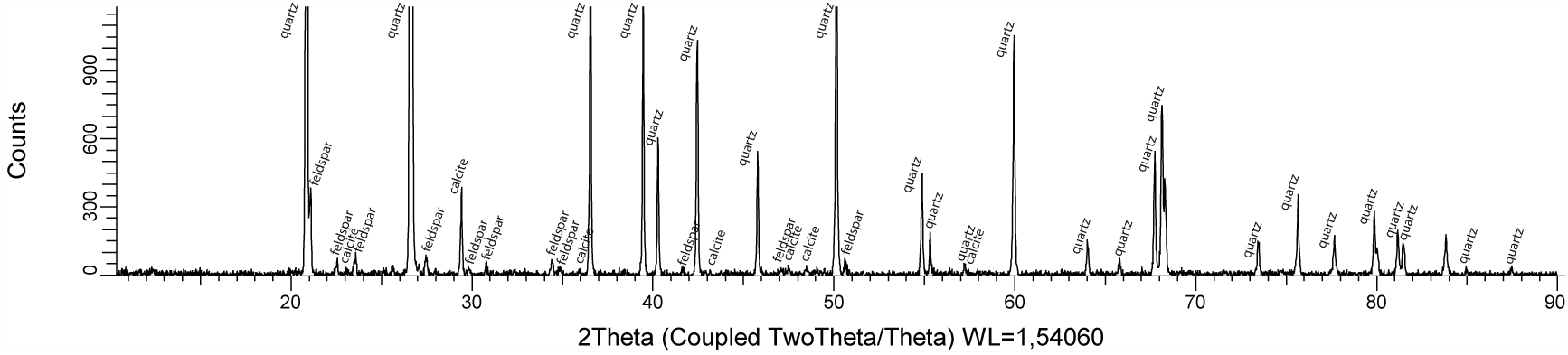
Representative XRD spectrum of consolidated sample (sample 18). Peaks for quartz, orthoclase feldspar and calcite are marked.

**Figure 9:**
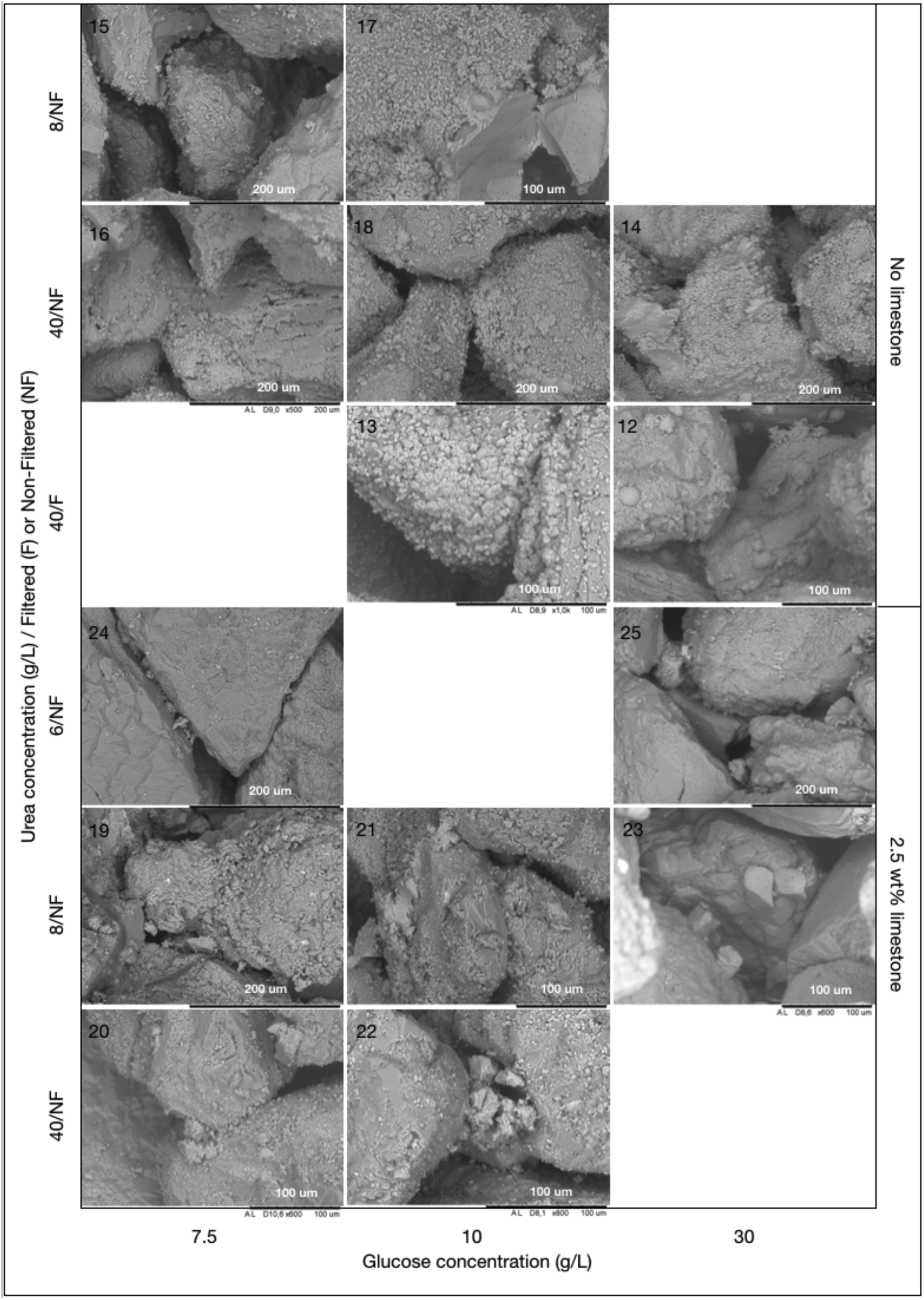
SEM observations of prototype samples 12-25. Sample numbers are shown in the top left corner of each image.

Samples 12-25 were tested mechanically and CaCO_3_ content measured (Table 2). They were also immersed in deionized water for at least 48 hours to see if the consolidated portions remained intact, which all did except sample 23. This suggests that this sample was consolidated by water-soluble by-products or organic material rather than calcium carbonate. For samples 23 and 25, the calculated CaCO_3_ content was negative, probably due to insufficient performance of the bottom filter (Figure 3), leading to substantial wash-out of limestone and bacterial CaCO_3_ precipitates into the outlet tube.

**Table 2:** Results from mechanical testing, measured CaCO_3_ content, and retained consolidation in water. These measurements were only done for samples 12-25.

| Sample nr | Compressive strength (MPa)<br>(3 duplicates) | $\text{CaCO}_3$ content (wt %) | Retained consolidation in water |
| --- | --- | --- | --- |
| 12 | -<br>-<br>0.25 (top), 0.37 (bottom) | 4.4<br>5.4<br>5.7 | yes |
| 13 | 0.29 (top), 0.26 (bottom)<br>0.30<br>0.24 | 4.4<br>4.9<br>3.2 | yes |
| 14 | -<br>- | 0.6<br>0.8 | yes |
|  | 0.95 (top), 1.22<br>(bottom) | 5.1 |  |
| 15 | -<br>-<br>0.72 | 1.4<br>2.3<br>2.4 | yes |
| 16 | 0.28<br>0.13<br>0.26 (top), 0.12<br>(bottom) | 1.7<br>3.4<br>2.9 | yes |
| 17 | -<br>-<br>0.24 | 4.5<br>3.9<br>5.2 | yes |
| 18 | 0.25 (top), 0.30<br>(bottom)<br>-<br>3.6 | 3.7<br>2.7<br>7.5 | yes |
| 19 | -<br>0.24<br>0.34 (top), 0.42<br>(bottom) | 3.7<br>2.8<br>2.7 | yes |
| 20 | 0.40 | 0.8 | yes |
|  | 0.29 | 1.5 |  |
|  | 0.15 | 0.9 |  |
| 21 | 0.06 | 2.1 | yes |
|  | - | 2.8 |  |
|  | 0.13 | 3.6 |  |
| 22 | - | 5.6 | yes |
|  | - | 2.0 |  |
|  | - | 3.9 |  |
| 23 | - | -1.9 | no |
|  | - | -1.4 |  |
|  | 0.72 | -1.7 |  |
| 24 | 0.18 | 2.6 | yes |
| 25 | 0.61 | -0.4 | yes |

In general, the mechanical strength and the CaCO_3_ content was highly variable, both between samples, between replicates and between different parts of the same sample. In addition, no significant correlation was observed between CaCO_3_ content and mechanical strength. These observations reflect the fact that consolidation in these samples was highly heterogeneous (Figure 7). The heterogeneous distribution of precipitated CaCO_3_ and thus mechanical strength is most likely influenced by the injection method, where air was introduced between each injection. This would favour the development of preferential flow paths through the porous material, with trapped air left in patches of the material during injections.

The uniaxial compressive strengths ranged from 0.15 to 1.22 MPa (one sample at 3.6 MPa) and the CaCO_3_ contents from 0.6 to 7.5 wt%. In a comparable study using dissolved eggshells as a calcium source, Choi *et al.* [24] reported compressive strengths in the range 0.3 to 0.5 MPa for CaCO_3_ contents between 5 and 8 %, which is consistent with our results. The same group achieved slightly higher values, 0.5 to 1.4 MPa, in another study where limestone was dissolved in acetic acid derived from pyrolysis of lignocellulosic biomass [16], for CaCO_3_ contents between 6 and 8 %. Acid concentrations in this study were in the range of 200 to 1200 mM, with the highest calcium concentration, 830 mM, produced for the lowest acid concentration. Uniaxial strengths as high as 40 MPa have been reported for MICP applications [25], showing that the method we present here is by no means optimised in terms of injection strategy, aggregate composition, and other processing parameters. However, the consolidation levels we achieved with these simple prototypes indicate that this is in fact a feasible consolidation process and a viable basis for further optimization in subsequent studies.

## 4. Discussion

### 4.1 A two-stage biocementation process as prerequisite for the ultimate goal of an integrated one-step process

The presented work describes and demonstrates a new concept for biocementation employing BioZEment as a binder. Bacterial metabolic processes are employed in two subsequent steps, first to obtain free Ca^2+^ and bicarbonate ions, and next to trigger recrystallization of CaCO_3_ for the consolidation of granular matter. The use of bacteria *in situ* has the potential to reduce costs by eliminating costly production steps such as purification of acid and/or enzymes. In addition, it may be a prerequisite to maintain liberated carbonate ions in solution and prevent irreversible loss of CO_2_ from the material. The new concept also holds promise towards a potential 1-step, *in situ* biocementation process. In this case, a carefully designed biological system may allow biologically catalysed dissolution and re-precipitation processes to be separated in time and thereby be carried out without the need for a separate dissolution reactor and multiple injections of reactant. In the current study, we have divided this ideal process scenario into two stages: (1) bacterially induced dissolution of CaCO_3_ and (2) bacterially induced CaCO_3_ re-crystallization. This separation allowed us to study the two processes independently, and enabled a proof-of-concept of consolidation with less stringent performance requirements for the two stages. Further essential fundamental studies and optimization of the process can be easier performed and results interpreted based on individual stages, prior to combining them into one efficient continuous process in the future.

### 4.2 The necessity for acid production at high initial pH and with a high content of solid limestone with optimal rate and maximum concentration

A slurry of powdered limestone (calcium carbonate) provides a high pH environment where only specially adapted bacterial strains can grow. As an additional constraint, a suitable strain must produce significant amounts of acid from glucose (preferably even homofermentatively) to enable limestone dissolution. Strain AP-004, selected for the present study based on acid production profile and ease of handling, is thereby just one example of potentially suitable strains for this purpose. It is not excluded that new isolates and/or optimization by genetic engineering are needed towards a potential future commercial application.

The rate of limestone dissolution *in situ* is proportional to the surface area of the solids in solution as well as the acid production rate, and a large surface area was targeted in the current scenario. This is in contrast to conventional fermentation processes for acid production on a given substrate where calcium carbonate is sometimes added continuously in small amounts as a pH buffer throughout the process [26]. In this case, the goal is to produce a high amount of acid and not to dissolve a high amount of calcium carbonate.

The acid concentrations currently obtained with strain AP-004 are low compared to other acid producing microbial systems. It should therefore be possible to optimise the microbial system, based on further development of AP-004 or isolation of other suitable strains, to achieve much higher concentrations of produced acid. Lactic acid fermentation by fungi, yeasts, and bacteria [27] are reported to yield up to 1035 mM lactic acid by using a glucose concentration of 150 g/L and at a pH around 6.40. Thereof, the bacterial production (*Lactobacillus sp.*) was reported to yield around 460 mM lactic acid. Acid yield well above 1 M (at pH 6.2) was also shown by Xu et al [28]. Our experiments showed that limited oxygen levels were necessary for effective acid production and that the acid production was better in larger batches with more efficient mixing. We expect that optimisation of the medium, including the glucose concentration, can result in higher production rates and final concentrations of acid and dissolved Ca^2+^. On the other hand, previous studies have shown that moderate concentrations of Ca^2+^ (in the range of 100-500 mM, compared with 1 M) yield the best quality of cementation, due to more homogeneous and slower growth of calcium carbonate crystals [29]. Further work is therefore required to find the optimal acid concentration and corresponding concentration of Ca^2+^ in this system.

### 4.3 The need for improvements in recrystallization and process configuration based on systems scale understanding

In the second stage of the BioZEment process, consolidation is achieved through the recrystallization of dissolved limestone on the surfaces of a granular material. At present, it is difficult to combine dissolution and recrystallization *in situ*. The repetitive injection process described above was developed and applied to achieve a first proof-of-concept. Consolidation based on the urease producing capabilities of *S. pasteurii* is fast, on the time scale of hours, while the dissolution process currently require several days. Details of the consolidation process are revealed by the microstructure of formed CaCO_3_ and their interactions with the granular media in the sample. We observed that sand grains indeed were connected and material consolidated by newly formed CaCO_3_ crystals. However, a deeper understanding of the recrystallization process in the complex mixture of sand, calcium carbonate crystals, cells of two bacteria strains, metabolic by-products and growth medium residues, will require a systems scale approach integrating both bacterial, geochemical, and mechanical aspects.

Future studies also need to address the effect of specific organic molecules present in solution on the reactivity of calcite surfaces. Organic molecules with hydrophilic functional groups have a tendency to bind to calcite surfaces [30], and the existence of such organic layers can significantly affect both dissolution and precipitation kinetics. In addition, some organic acids may lead to formation of poorly soluble precipitates that can form passivating layers on the calcite surfaces, hindering further dissolution.

In the precipitation step, urease is not the only factor that influences the system. Other studies have shown that the enzyme carbonic anhydrase, ubiquitous in many microorganisms and catalysing the interconversion of HCO_3^-^_ and CO_2(aq)_ [31] can play an important role in complex carbonate systems such as the BioZEment process. In addition, extracellular polymeric substances (EPS) may influence the MICP process by trapping Ca^2+^ ions and serving as nucleation sites [31], and the composition of the EPS may have a large effect on the growth and morphology of precipitated calcium carbonate crystals [32].

Compressive strength and stability was measured for several groups of prototype samples. The uniaxial compressive strength ranged from 0.15 to 1.22 MPa (one sample at 3.6 MPa) with CaCO_3_ contents between 0.6 and 7.5 wt%. In previously reported studies of microbial cemented sand, uniaxial compressive strength ranged between 0.15 to 34 MPa [29]. Even for similar amounts of precipitated CaCO_3_, the strength of the consolidated sand depends on the precipitation mechanism and the type and packing of aggregate material.

We also found that the compressive strength of our samples were highly variable and inhomogeneous. This was probably at least partly due to the sample preparation procedure, where sand was poured loosely into moulds and pressed down by a rubber plug. This did most likely not lead to the maximum possible level of compaction and could also have created regions of loose material inside the moulds. Another reason for variable consolidation is the injection strategy itself, where the supply of reagents to parts of the material could have been reduced by incomplete wetting or by clogging of flow paths due to precipitation. It is well known that injection methods are prone to non-uniform distribution of precipitated material [29]. One approach to counter this problem is to use staged injections, where bacteria are first injected at a slow rate, followed by a pause to ensure that the bacteria become fully attached to the sand grains. Then, the reagent is injected at a higher rate in order to be distributed in the medium before the onset of the precipitation process. Another promising approach is the surface percolation method, where a bacterial solution and reagent solution are sprayed alternatingly onto the sand surface and allowed to percolate down by gravity. This has been shown to yield fully cemented soil up to 2 m depth [29], but to work best for coarse-grained sands and soils.

### 4.4 Provisional cost analysis and Life Cycle Assessment identify key drivers towards economic feasibility, sustained public acceptance, and reduced climate impact

Initial Techno-Economic Analysis (TEA) and Life Cycle Assessment (LCA) were performed in order to identify at an early stage bottlenecks in the proposed process **(Myhr et al, submitted)**. Although the knowledge about the final process components is limited, we believe that such aspects need to be considered early in the product development in order direct the ongoing research toward key challenges. TEA and LCA revealed that both production time and water use directly influenced cost, and hence market penetration potential, and the climate impact of the material produced with BioZEment. Production time and water use may be reduced by achieving a higher amount of Ca^2+^ in the injected solution, through enabling the bacteria to produce acids up to higher concentrations and ensuring that the dissolution of calcium carbonate proceeds to equilibrium. This would allow the production to take place using fewer injections of reagent and hence a shorter total production time. In addition, a higher rate of acid production may reduce the time needed in the dissolution step of the production.

Arguably, public acceptance issues more than ever have become a primary concern in innovation efforts. In general, there are concerns over innovations based on emerging technologies [33]. This has definitely included biotechnological innovations. In particular, the construction community is not used to dealing with biological processes and concerns of pathogenicity and other health issues will need to be taken into account as these processes are moving closer to market [3].

## 5. Conclusion

We have presented a novel method for biocementation using BioZEment, in which calcium carbonate, in the form of powdered limestone, is dissolved and subsequently recrystallized through microbial mediated processes. A solid material has been produced when calcium carbonate precipitated on the solid surfaces of aggregate grains and bound them together in a dense network. The use of cheap, abundantly available raw materials such as limestone and glucose (which could be replaced with other sustainable, not food-competing sugar sources in future applications) provides an avenue for material production with low energy demand and environmental footprint. Our results also indicate the possibility of a fully integrated one-step process where bacterial dissolution and precipitation both take place in a densely packed granular system of limestone and sand, but separated in time. In conclusion, the current study represents uncharted territory in industrial biotechnology with multiple remaining challenges both within experimentation and interpretation of results that need to be addressed and solved in subsequent and currently ongoing activities. However, the presented proof-of-concept provides a promising and motivating basis for this, on the way to a comprehensive understanding that will ultimately enable directing the involved molecular processes towards a competitive and more sustainable concrete-like building material in the future.

## Acknowledgements

The authors would like to thank Tone Haugen, Kari Hjelen, and Kristin Reine at SINTEF Industry for their contributions in strain isolation and characterization. The remaining members of the BioZEment team, Anita Borch and Frida Røyne, are acknowledged for their contributions in fruitful and inspiring discussions.

